# Structure of mycobacterial respiratory Complex I

**DOI:** 10.1101/2022.10.04.510895

**Authors:** Yingke Liang, Alicia Plourde, Stephanie A. Bueler, Jun Liu, Peter Brzezinski, Siavash Vahidi, John L. Rubinstein

## Abstract

Oxidative phosphorylation, the combined activity of the electron transport chain (ETC) and adenosine triphosphate synthase, has emerged as a valuable target for the treatment of infection by *Mycobacterium tuberculosis* and other mycobacteria. The mycobacterial ETC is highly branched with multiple dehydrogenases transferring electrons to a membrane-bound pool of menaquinone and multiple oxidases transferring electrons from the pool. The proton-pumping type I nicotinamide adenine dinucleotide (NADH) dehydrogenase (Complex I) is found at low abundance in the plasma membranes of mycobacteria in typical *in vitro* culture conditions and is often considered dispensable. We found that growth of *Mycobacterium smegmatis* in carbon-limited conditions greatly increased the abundance of Complex I and allowed isolation of a rotenone-sensitive preparation of the enzyme. Determination of the structure of the complex by cryoEM revealed the “orphan” two-component response regulator protein MSMEG_2064 as a subunit of the assembly. MSMEG_2064 in the complex occupies a site similar to the proposed redox sensing subunit NDUFA9 in eukaryotic Complex I. An apparent purine nucleoside triphosphate within the NuoG subunit resembles the GTP-derived molybdenum cofactor in homologous formate dehydrogenase enzymes. The membrane region of the complex binds acyl phosphatidylinositol dimannoside, a characteristic three-tailed lipid from the mycobacterial membrane. The structure also shows menaquinone, which is preferentially used over ubiquinone by gram-positive bacteria, in two different positions along the quinone channel and suggests that menaquinone interacts more extensively than ubiquinone with a key catalytic histidine residue in the enzyme.

## Introduction

The bacterial genus *Mycobacterium* includes human pathogens such as *M. tuberculosis, M. leprae, M. abscessus*, and *M. avium*, as well as non-pathogenic species such as *M. smegmatis*. These bacteria are considered obligate aerobes due to their dependence on oxidative phosphorylation for generating adenosine triphosphate (ATP) (1). In oxidative phosphorylation, the electron transport chain (ETC) couples the oxidation of metabolites to the transport of protons out of the cytoplasm across the mycobacterial inner membrane, establishing an electrochemical protonmotive force (PMF). The ATP synthase allows protons to flow back into the cytoplasm, using the free energy released in the process for the generation of ATP. The discovery and subsequent approval of the tuberculosis drug bedaquiline, which inhibits mycobacterial ATP synthase, identified oxidative phosphorylation as a valuable target for new antibiotics for mycobacteria (2, 3). Subsequently, multiple inhibitors of the mycobacterial ETC have been found to have antimycobacterial activity (3–7).

Despite requiring oxygen for growth, mycobacteria can survive for an extended period in low oxygen environments, enabling pathogens such as *M. tuberculosis* to persist as latent infections within granulomas in the lungs (8, 9). Response to hypoxia is regulated by two-component signalling systems such as the DosS/R regulon (10) that modulates mycobacterial oxidative phosphorylation (11). Metabolic flexibility in mycobacteria (1, 12), including resistance to hypoxia, is supported by a highly-branched ETC. Unlike the linear ETCs in mammalian mitochondria and many bacteria, in mycobacteria multiple oxidoreductases accept electrons from diverse sources and transfer them to the lipid-soluble electron carrier menaquinone (MQ), reducing it to menaquinol (MQH_2_) (13). Electrons from MQH_2_ then exit the ETC through multiple terminal oxidases (14). These electron entry and electron exit branches can include catalysis of the same reaction by two or more apparently redundant enzymes, which complicates inhibition of the mycobacterial ETC. Previously-determined structures from the mycobacterial ETC include the succinate dehydrogenases Sdh1 (15) and Sdh2 (16) (also known as respiratory Complex II), the cytochrome *aa*_*3*_*-bcc* terminal oxidase (17–20) (also known as the Complex III_2_IV_2_ respiratory supercomplex), and the cytochrome *bd* terminal oxidase (21, 22).

One of the main points for electron entry into the ETC is from reduced nicotinamide adenine dinucleotide (NADH) produced by the Krebs cycle, glycolysis, and fatty acid oxidation. Electrons from NADH enter the mycobacterial ETC through two different NADH:menaquinone oxidoreductases (also known as NADH dehydrogenases), either of which is sufficient for mycobacterial survival *in vivo* (23). Type II NADH dehydrogenases, such as the Ndh and NdhA proteins in *M. tuberculosis* and the Ndh protein in *M. smegmatis* (24), are small dimeric peripheral membrane proteins that do not contribute to the PMF. Type I NADH dehydrogenases, also known as respiratory Complex I, are large L-shaped multimeric membrane-embedded assemblies that translocate four protons for each NADH molecule oxidized (25). Structures have been determined for intact Complex I from the mitochondria of mammals (26–28), vascular plants (29), yeast (30), and ciliates (31). However, intact Complex I structures from bacteria have been determined only for *Thermus thermophilus* (32) and *Escherichia coli* (33). The core Complex I structure consists of 14 subunits, with 31 supernumerary subunits in mammalian mitochondria (28, 34), and total molecular masses ranging from ~500 kDa in prokaryotes to >1 MDa in eukaryotes. Within Complex I, electrons from NADH are accepted by a flavin mononucleotide (FMN) cofactor before sequential transfer through eight iron-sulfur clusters (N1a, N1b, N2, N3, N4, N5, N6a, and N6b) to the membrane-embedded quinone, either ubiquinone (UQ), which is the sole quinone species in mitochondria (35), or MQ. An additional iron-sulfur cluster in Complex I found only in some prokaryotes, N7, has been proposed to help maintain the enzyme’s structure (36).

All mycobacteria other than *M. leprae* possess the *nuoAN* operon that encodes Complex I, but the enzyme has been described as dispensable in both *M. tuberculosis* and *M. smegmatis* (37–39). However, Complex I has also been found to be important for virulence in *M. tuberculosis* (24, 40) and transcription of the *nuoAN* operon differs by more than 20-fold in *M. smegmatis* depending on the carbohydrate source in the bacterial growth medium (12). Knockout of the genes for type II NADH dehydrogenases in *M. tuberculosis* makes the bacterium highly sensitive to the Complex I inhibitor rotenone, while knockout of all NADH dehydrogenase genes renders the bacterium non-viable (23, 24).

We found that although Complex I protein levels were extremely low in *M. smegmatis* cultured under standard laboratory conditions, the complex was easily detected in bacteria grown in media that was not supplemented with carbohydrates. Addition of glucose to the growth medium decreased Complex I levels more than succinate or glycerol and anaerobic growth prevented detection of Complex I regardless of the composition of the medium. Growth of *M. smegmatis* in carbon-limited conditions allowed purification of Complex I, which displayed rotenone-sensitive NADH:quinone oxidoreductase activity. CryoEM of this protein preparation enabled calculation of a map of the complex at 2.6 Å resolution. This map, combined with mass spectrometry, revealed that the two-component response regulator MSMEG_2064, for which the cognate kinase is currently unknown, is a subunit of the complex. An additional density adjacent to iron-sulfur cluster N7 fits a purine nucleoside triphosphate molecule, supporting the homology between the NuoG subunit and formate dehydrogenases that employ a GTP-derived molybdenum cofactor in the same position. The abundant mycobacterial lipid acyl phosphatidylinositol dimannoside was found in the structure at the interface of subunits NuoL and NuoM. Finally, MQ was resolved in two positions within the quinone channel, suggesting that the overall mechanism of quinone binding is conserved between UQ and MQ. However, interactions between MQ and a key histidine residue in the quinone binding pocket are more extensive for MQ than UQ, which may explain how Complex I NADH:menaquinone oxidoreductase activity can occur at rates comparable to NADH:ubiquinone oxidoreductase activity despite the weaker oxidizing potential of MQ (41).

## Results

### Active mycobacterial Complex I can be isolated from M. smegmatis in carbon-limited growth

To enable quantification and isolation of Complex I from *M. smegmatis*, a sequence encoding a C-terminal 3×FLAG tag was incorporated into the bacterial genome 3’ of the gene for the soluble Complex I subunit NuoC. The bacteria were cultured in a variety of different conditions and NuoC-3×FLAG was detected by western blotting (Fig. 1A, *upper*). All conditions included Middlebrook 7H9 broth with tween-80 detergent, which reduces aggregation of cells and can be metabolized by the bacterium (42). The conditions differed in the presence or absence of albumin, sodium chloride, D-glucose, succinate, and glycerol. In some conditions, Wayne’s model of hypoxia was employed (43). Coomassie staining of membranes after western transfer (Fig. 1A, *lower*) shows numerous changes in the protein composition of the bacteria under these conditions. Consequently, loading was controlled by adding a total of 4 μg of protein to each lane of the SDS-PAGE gel rather than blotting against a single control protein. Comparison of anti-FLAG signal for the different conditions shows that NuoC is most abundant with little or no carbohydrate added to the growth medium (Fig. 1A, *upper*, lanes 2, 3, 5, 6, and 7). At carbohydrate concentrations typical for *in vitro* growth (0.2% or 11 mM glucose), NuoC is less abundant (Fig. 1A, *upper*, lanes 1 and 4). In hypoxic conditions the NuoC protein cannot be detected (Fig. 1A, *upper*, lanes 8, 9, and 10). In contrast, protein levels for the type II NADH dehydrogenase Ndh did not vary as strongly between growth conditions (Fig. 1B). This result suggests that variability in the amount of Complex I contributes to the flexibility of the mycobacterial NADH dehydrogenase system. We next constructed a *M. smegmatis* strain with a deletion of the *nuoF* gene. This gene encodes the NuoF subunit that binds the FMN moiety, which accepts electrons from NADH and is therefore essential for Complex I activity. However, the Δ*nuoF* strain did not display a noticeable growth defect in conditions where Complex I is expressed, suggesting that even in conditions where Complex I is expressed it is not essential for growth.

**Figure 1:**
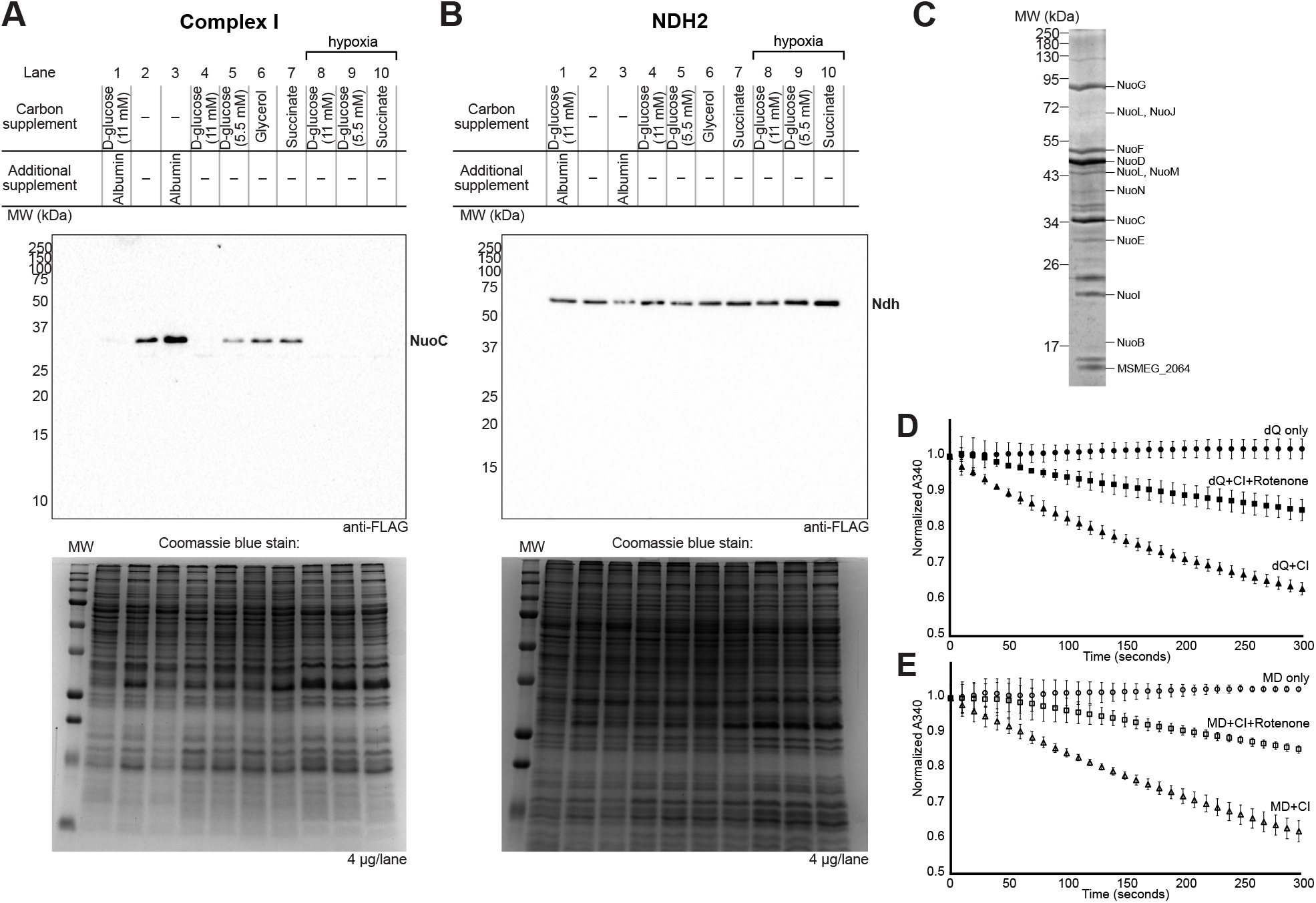
Biochemical characterization of mycobacterial Complex I. **A**, Western blot of 3×FLAG-tagged Complex I subunit NuoC under different culture conditions (*upper*) with a Coomassie blue stained membrane showing total loaded protein (*lower*). MW, molecular weight. **B**, Western blot of 3×FLAG-tagged NDH2 under different culture conditions (*upper*) with a Coomassie blue stained membrane showing total loaded protein (*lower*). **C**, Coomassie blue-stained SDS-polyacrylamide gel of purified *M. smegmatis* Complex I. **D**, Representative Complex I NADH:decylubiquinone (dQ) oxidoreductase activity with and without 50 mM rotenone. **E**, Representative Complex I NADH:menadione (MD) oxidoreductase activity with and without 50 mM rotenone.

To determine if the Complex I that is expressed is active, *M. smegmatis* was grown at a large scale in 7H9 medium supplemented with 11 mM sodium succinate. This growth condition is a compromise that enables production of sufficient biomass for protein purification while still inducing Complex I expression comparable to when media is supplemented with albumin alone. *M. smegmatis* membranes were isolated and solubilized with dodecyl-β-D-maltoside (DDM) and Complex I was isolated with anti-FLAG affinity chromatography and gel filtration chromatography. Screening of Complex I stability by gel filtration chromatography showed an optimum when buffers were adjusted to pH 6.0, similar to Complex I from *E. coli* and *T. thermophilus* (32, 33), and when the 3×FLAG tag was attached at the C terminus of the NuoH subunit. Polyacrylamide gel electrophoresis with the resulting protein preparation appeared to show all the expected subunits of Complex I except the small membrane subunit NuoA (Fig. 1C). This preparation was assayed for NADH:quinone oxidoreductase activity using either decylubiquinone (dQ, Fig. 1D and Fig. S1A, *triangles*) or menadione (MD, Fig. 1E and Fig. S1B, *triangles*) as the electron acceptor. With either quinone species, the enzyme had a mean specific activity of 5 μmol/min/mg (s.d ± 0.8 μmol/min/mg for decylubiquinone and ± 0.6 μmol/min/mg for menadione, n=6 independent assays from two separate batches of protein for each quinone). The enzyme does not display the de-active to active transition seen with Complex I from vertebrates and fungi (44), which is characterized by an initial delay in NADH oxidation on addition of the substrates to the assay (Fig. 1D and 1E, *triangles*). Rotenone, a competitive Complex I inhibitor that blocks the quinone binding site (45), decreased activity of the enzyme by ~50% at a concentration of 50 mM (Fig. 1D and 1E, *squares*, n=3 independent assays for each quinone). Interestingly, 150 mM rotenone was able to decrease NADH:decylubiquinone oxidoreductase activity by ~90% (Figure S1A, *squares*) but NADH:menadione oxidoreductase activity by only ~50% (Figure S1B, *squares*). However, neither decylubiquinone nor menadione are the *in vivo* electron acceptor for mycobacterial Complex I. The specific activity of the mycobacterial enzyme is similar to the ~5 μmol min^−1^ mg^−1^ activity of *E. coli* Complex I in lipid nanodiscs (33), although substantially less than the 15 to 22 μmol min^−1^ mg^−1^ activity of detergent-solubilized Complex I from *Paracoccus denitrificans* (46, 47) or the ~40 μmol min^-1^ mg^-1^ activity of *T. thermophilus* Complex I (48).

### Mycobacterial Complex I resembles the enzyme from other bacteria

CryoEM of purified Complex I allowed for calculation of two-dimensional (2D) class average images that suggest that the enzyme is highly flexible (Fig. S2A). *Ab initio* three-dimensional (3D) map calculation and map refinement revealed the structure at an overall nominal resolution of 2.6 Å (Fig. 2A, *top*, Fig. S2B-D, Table S1), with local resolutions ranging from 2.6 to 4.5 Å (Fig. S2E). This map enabled construction of an atomic model for 96% of the residues in the complex (Fig. 2A, *bottom*, Table S2, Fig S3). The structure shows that the overall architecture of mycobacterial Complex I resembles Complex I from other bacteria (32, 33). Like other Complex I structures, the membrane region consists of three transporter-like subunits (NuoL, NuoM, and NuoN) with NuoA, NuoJ, NuoK, and NuoH forming a possible fourth transporter module (30, 32). Each of these transmembrane modules contains polar residues within their core (Fig. S4A), which have previously been associated with proton transport (49). Three dimensional variability analysis (3DVA) (50) (Video 1 and 2) indicates conformational flexibility in the cryoEM dataset consistent with the apparent flexibility in the 2D class averages and conformational changes seen in previous cryoEM of Complex I (48, 28, 33). Two different modes of flexibility were observed: twisting of the peripheral arm and the membrane region relative to each other (Video 1), and a clamshell-like widening and narrowing of the angle between the two regions (Video 2). This variability highlights the dynamic nature of Complex I even in the absence of substrate turnover.

**Figure 2:**
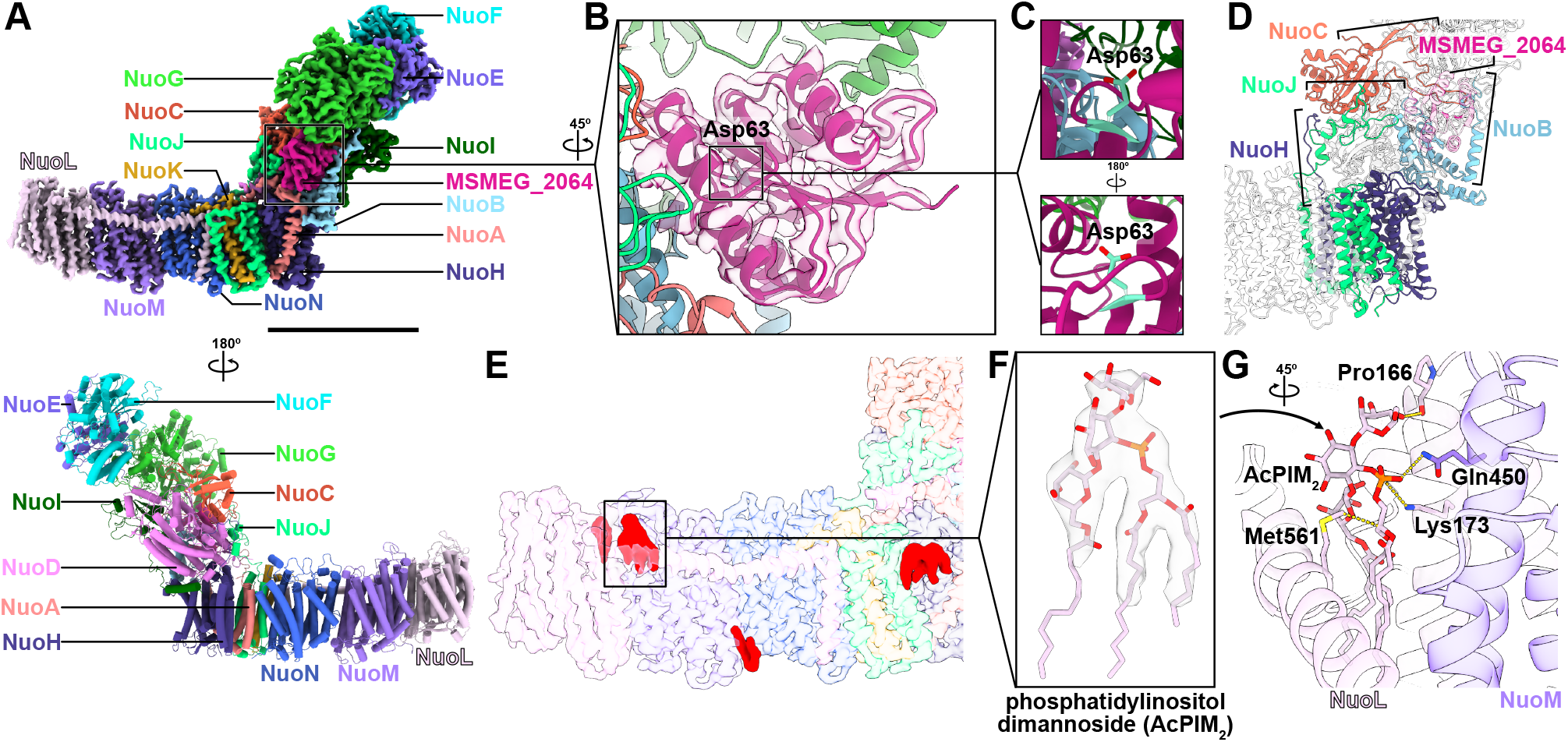
Structure of mycobacterial Complex I. **A**, CryoEM map of *M. smegmatis* Complex I (*top*) and corresponding atomic model (*bottom*) with subunits labelled. Scale bar, 100 Å. **B**, Model for MSMEG_2064 in cryoEM density. Inset indicates position of phosphorylation target Asp63. **C**, Asp63 is occluded when MSMEG_2064 is bound to Complex I. **D**, Atypical C-terminal extensions of NuoB, NuoC, NuoH, and NuoJ subunits. MSMEG_2064 is indicated in pink. **E**, Lipid densities in membrane region (*red*) from the cryoEM map. Inset indicates position of acyl phosphatidylinositol dimannoside (AcPIM_2_). **F**, Close-up view of AcPIM_2_ density with fitted atomic model. **G**, Main interactions between AcPIM_2_ with NuoM and NuoL.

### The two-component response regulator MSMEG_2064 is a subunit of mycobacterial Complex I

The mycobacterial Complex I structure shows the 14 core subunits, NuoA to NuoN. Density corresponding to an unidentified subunit (Fig. 2B, *semi-transparent pink surface*) is found adjacent to the NuoB subunit. In addition to the known Complex I subunits, mass spectrometry of the protein preparation detected MSMEG_2064 as a component of the sample (Table S4). MSMEG_2064 is found in the same operon as other Complex I genes 5′ of *nuoA* and was proposed to be a constituent of mycobacterial Complex I (12). Furthermore, recent crosslinking experiments suggested that MSMEG_2064 associates with NuoD and deletion of the MSMEG_2064 gene affects expression of some electron transport chain components (51). The structure of MSMEG_2064 predicted by AlphaFold (52) fits into the unidentified density with high fidelity (Fig. 2B, *pink ribbons*). In the structure, MSMEG_2064 forms extensive contacts with NuoB, as well as contacts with NuoA, NuoC, and NuoI, but not the NuoD subunit that it was proposed to contact from crosslinking.

MSMEG_2064 is a two-component response regulator protein that is considered to be an “orphan” because its cognate kinase has not been identified (51, 53). Sequence comparison and structural alignment with the PDBeFold server (54, 55) showed similar folds but low sequence identity (<30%) between MSMEG_2064 and other two-component response proteins with known structures. Cα root mean squared deviation (RMSD) for backbone structures was lowest between MSMEG_2064 and the *Caulobacter crescentus* two-component response regulator protein DivK (Table S3, Fig S4B). In response to a checkpoint signal late in the cell cycle, DivK is phosphorylated on Asp53 to initiate a phosphorelay cascade that ultimately leads to upregulation of genes involved in motility (56, 57). The homologous Asp63 residue in MSMEG_2064 is buried against NuoB in the structure (Fig. 2C), suggesting that it cannot be phosphorylated when MSMEG_2064 is bound to Complex I. To investigate the function of MSMEG_2064 in mycobacteria, we constructed a *M. smegmatis* strain bearing a 3×FLAG tag on NuoH and with the gene for MSMEG_2064 deleted. However, intact Complex I could not be purified from this strain, with affinity chromatography isolating just the membrane region of the complex (Fig. S4C). This observation suggests that MSMEG_2064 contributes to the assembly or structural integrity of mycobacterial Complex I.

### Atypical subunit extensions secure the peripheral arm to the membrane region

The subunits NuoB, NuoC, NuoH, and NuoJ in mycobacterial Complex I have extended sequences at their N or C termini compared to the corresponding subunits in Complex I from other species (Fig. 2D, *black brackets*). The C termini of NuoH and NuoJ extend from the membrane region to the peripheral arm suggesting that they stabilize the connection between these two regions of the complex (Fig. 2D, *green* and *dark blue ribbons*). Extensions bridging the peripheral arm and membrane region are not found in other bacterial Complex I structures (Gutiérrez-Fernández et al., 2020; Kolata and Efremov, 2021). While the C terminus of NuoJ is well resolved with clear secondary structure, the C-terminal sequence of NuoH displays weak density, suggesting that it is flexible. This region (residues 375-395) was modelled at low resolution, which could result some inaccuracy. A C-terminal extension from NuoB (residues 156-184) forms an α helix (Fig. 2D, *light blue ribbons*) that extends from the peripheral arm to bind MSMEG_2064 (Fig. 2D, *transparent pink ribbons*) and may be involved in its function. Finally, the N terminus of mycobacterial NuoC is highly extended compared to homologues and forms contacts with the NuoG subunit (Fig. 2D, *orange ribbons*). This NuoC extension may stabilize the NuoEFG module, which has been found to exist as a separate subcomplex during assembly of *E. coli* Complex I (58). These extensions in mycobacterial Complex I subunits highlight the diversity of Complex I structures even within bacteria.

### The mycobacterial phospholipid AcPIM_2_ binds the NuoM-NuoL interface

Several densities corresponding to lipids are found in the cryoEM map (Fig. 2E, *red surfaces*). While most of these densities could not be assigned as a specific lipid, a distinctive and well-resolved lipid is found at the interface between NuoM and NuoL (Fig. 2F). The resolution of this density and its distinctive shape allowed it to be assigned as the three-tailed lipid acyl phosphatidylinositol dimannoside (AcPIM_2_). AcPIM_2_, along with the related four-tailed lipid diacyl phosphatidylinositol dimannoside (Ac_2_PIM_2_), is unique to mycobacteria and highly abundant in the inner membrane (59–61). AcPIM_2_ comprises ~10% of the *M. smegmatis* inner membrane (62), with Ac_2_PIM_2_ proposed to comprise almost all of the inner leaflet of the inner membrane. In the structure, the phosphatidylinositol headgroup of AcPIM_2_ is positioned to form contacts with Pro166, Lys 173, and Met561 of NuoL, and Gln450 of NuoM (Fig. 2G, *dashed yellow lines*).

Synthetic phosphatidylinositol mannoside (PIM) has previously been co-crystallized with a mammalian protein (63), but observation of AcPIM_2_ binding at the NuoM–NuoL interface is the first time endogenous PIM has been resolved clearly in a protein structure. The unusual lipid composition of the mycobacterial inner membrane has been suggested to promote stability of the inner membrane and resistance to drugs (59). The binding of the lower abundance AcPIM_2_ to the NuoM-NuoL interface suggests that it may stabilize NuoL during movement as the distal portion of the membrane region appears to be particularly flexible (Video 1 and 2). Multiple lipids are found in cryoEM maps of eukaryotic Complex I (26) and have also been proposed to play a role in conformational dynamics in the complex (64).

### Iron-sulfur clusters electronically connect FMN to MQ in the peripheral arm

Ten redox centers form the electron transfer pathway within the peripheral arm of mycobacterial Complex I (Fig. 3A and B): one FMN, two 2Fe–2S iron sulfur clusters (N1a and N1b), and seven 4Fe–4S iron sulfur clusters (N2, N3, N4, N5, N6a, N6b, and N7). Density corresponding to a quinone molecule is found at the base of the peripheral arm where it joins the membrane region (Fig. 3A and B, *labelled as ‘Q’*). The nine iron-sulfur clusters are found in other prokaryotic Complex I structures, including the N7 cluster that is unique to some prokaryotes. However, bound quinone was not observed in previous structures of bacterial Complex I (32, 33, 48) and is discussed in more detail below. Of the nine iron sulfur clusters, seven (N1a, N1b, N2, N3, N4, N6a, and N7) are coordinated by four cysteine residues, as seen in Complex I from other organisms. Cluster N5 is coordinated by three cysteines and one histidine (Fig. 3C), which is a characteristic feature of this cluster (65). Histidine coordination of iron-sulfur clusters has been suggested to tune their midpoint potentials, as seen when comparing dihistidine–dicysteine coordination of Rieske-type clusters to typical tetracysteine coordinated 2Fe–2S clusters (65). In addition to cluster N5, cluster N6b in the mycobacterial complex interacts with His42 of NuoI within coordination distance (Fig. 3D). This interaction between cluster N6b and a histidine residue has not been observed in Complex I from other species and could suggest a difference in its midpoint potential compared to other species.

**Figure 3:**
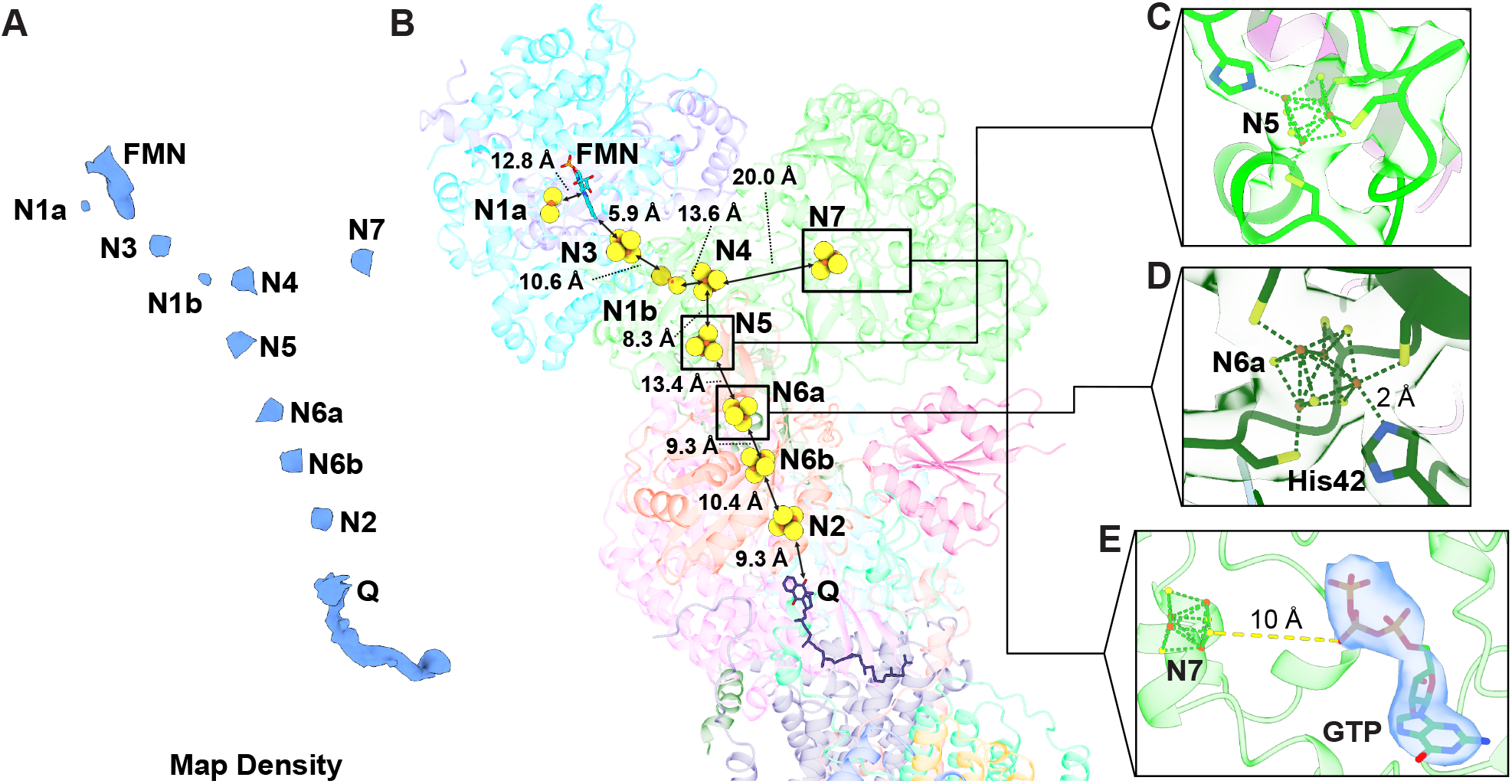
The peripheral arm cofactors of mycobacterial Complex I. **A**, CryoEM density from redox centres in peripheral arm of mycobacterial Complex I. **B**, The complete redox network of *M. smegmatis* Complex I consists of a flavin mononucleotide (FMN), nine iron-sulfur clusters (N1a–N7), and a menaquinone-9 (labelled *‘Q’*). Edge-to-edge distances between adjacent redox centres are labelled. **C**, Coordination of cluster N5. **D**, Coordination of cluster N6a. **E**, The unassigned purine nucleoside triphosphate density, modelled as a GTP, found ~10 Å from cluster N7 in the NuoG subunit.

During electron transfer, NADH donates two electrons to the FMN bound non-covalently within NuoF, reducing it to FMNH_2_. The two electrons travel one-at-a-time from FMN through clusters N3, N1b, N4, N5, N6a, and N2, before ultimately reducing the MQ that is bound adjacent to N2. The distances between these cofactors (Fig. 3B) are sufficiently small to allow rapid electron transfer. Cluster N1a has been proposed to mediate transfer of the second electron from FMNH_2_ into the chain of iron-sulfur clusters in *E. coli* Complex I (66). However, cluster N1a could not be reduced by NADH in Complex I from species other than *E. coli* and its role remains contested (67). Cluster N7, which is found only in some prokaryotes, is closest to cluster N4 but the ~20 Å edge-to-edge distance between these two clusters is too far for rapid electron transfer (68, 69). Therefore, cluster N7 appears electronically disconnected from the electron transfer pathway between NADH and MQ, as it is in both *T. thermophilus* and *E. coli* (32, 33, 70).

### NuoG contains a GTP-like moiety similar to the homologous formate dehydrogenase

In addition to the expected cofactors in Complex I, an unexpected cofactor density is found ~10 Å from cluster N7 in NuoG (Fig. 3E, *semi-transparent blue surface*). This density corresponds in size and shape to a purine nucleoside triphosphate molecule, which we modelled as a guanosine triphosphate (GTP). Prokaryotic NuoG is homologous with metal-dependent formate dehydrogenases, which contain the GTP-derived molybdenum cofactor bis-molybdopterin guanine dinucleotide (Mo-bisMGD) (71, 72). When the atomic model for NuoG is overlayed with structures of the *E. coli* formate dehydrogenase FdhF (73) or the *Rhodobacter capsulatus* FdsA (74), the position of the modelled GTP corresponds to the guanosine moiety of the Mo-bisMGD cofactor that receives electrons from formate in those enzymes (Fig. S5A and B). Furthermore, the position of iron-sulfur cluster N7 in NuoG corresponds to the position of the iron-sulfur cluster proximal to the Mo-bisMGD in both formate dehydrogenase structures. However, assay of Complex I purified from *M. smegmatis* grown in medium supplemented with molybdenum did not detect formate:NAD^+^ oxidoreductase activity (Fig. S5C).

### Menaquinone binds the catalytic site more tightly than ubiquinone

Mycobacteria use MQ as an intermediate electron carrier rather than the UQ used in mitochondria (13, 75). MQ has a more negative midpoint potential than UQ (76, 77), which makes MQ a weaker oxidizing agent than UQ and MQH_2_ a stronger reducing agent than UQH_2_. The primary MQ species in mycobacteria is menaquinone-9 (MQ9) (75), which possesses a chain of nine isoprene units and a 2-methyl-1,4-napthaquinone ring that can carry two protons upon reduction of the ketone groups to hydroxyl groups (Fig. 4A). In the cryoEM map, the quinone binding channel is found at the interface of the NuoB, NuoD, and NuoH subunits (Fig. 4B, *solid black box*). The density for MQ in this channel possesses a noticeable bulge in the middle (Fig. 4B, *blue surface within dotted box*) that suggested overlap of two MQ9 molecules, as has been seen in previous respiratory complex structures (78). 3DVA with a mask including the quinone binding channel allowed separation of this density into two distinct densities (Fig. 4C and D, *semi-transparent blue surfaces*). Each of the separated densities accommodates a MQ9 molecule in either a fully-inserted position ~10 Å from iron-sulfur cluster N2 (Fig 4C, *yellow atomic model*), or semi-inserted position ~25 Å from cluster N2 (Fig. 4D, *yellow atomic model*), although density corresponding to the isoprene tail of MQ9 is not resolved from the micelle in the semi-inserted position (Fig. 4D, *blue surface*). The two MQ binding positions are similar to those observed with endogenous UQ in eukaryotic Complex I (29, 30, 79, 80), supporting a conserved quinone binding mechanism in Complex I.

**Figure 4:**
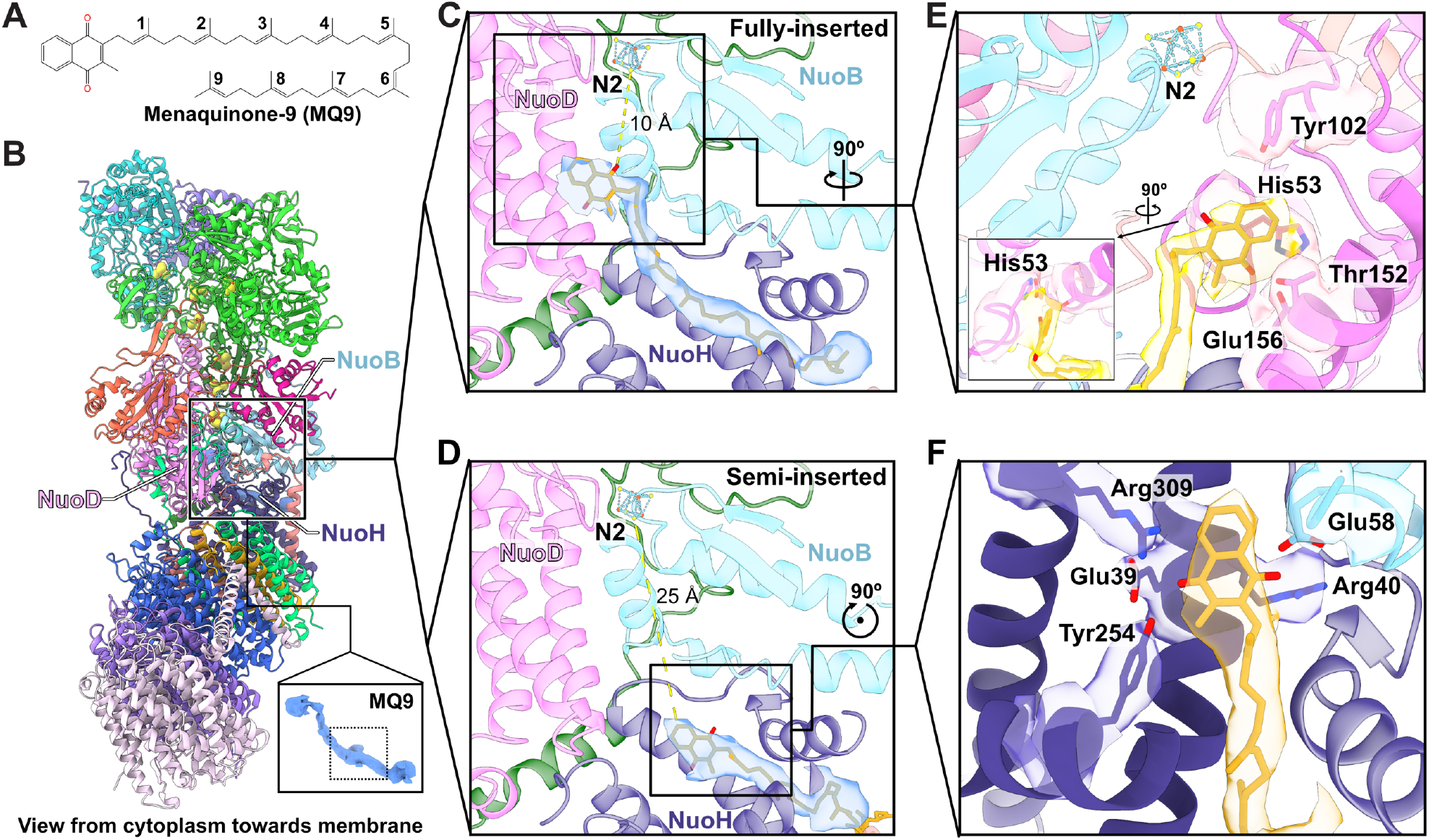
Menaquinone binds mycobacterial Complex I in multiple positions. **A**, Structure of menaquinone-9 (MQ9) with isoprene units numbered. **B**, The quinone binding site at the interface of NuoH, NuoB, and NuoD with density accommodating a full-length MQ9. Inset shows the consensus quinone density with a bulge highlighted by dotted square. **C**, CryoEM map and atomic model of MQ9 in the fully-inserted position. **D**, CryoEM map and atomic model of MQ9 in the semi-inserted position **E**, Sidechain interactions of mycobacterial Complex I with MQ9 in the fully-inserted position. Inset shows a closeup view of the interaction between MQ9 and His53. **F**, Sidechain interactions of mycobacterial Complex I with MQ9 in the semi-inserted position.

In the fully-inserted position, the naphthoquinone head of MQ is ~3 Å from Tyr102 of NuoD, which along with His53 of NuoD plays a key role in quinone binding and reduction (81, 82). The more elongated naphthoquinone head group of MQ allows it to form more extensive contact with His53 compared to UQ and the homologous His92 of NDUFS2 in porcine Complex I due to the smaller distance between MQ and His53 (Fig. S6). The smaller distance between the aromatic naphthoquinone group of MQ and His53 of mycobacterial Complex I allows for π-π stacking, further strengthening the interaction. A stronger interaction of His53 with MQ compared to UQ may kinetically compensate for the lower midpoint potential of MQ to allow similar turnover rates regardless of the quinone species (41). In the semi-inserted position, the naphthoquinone interacts with Glu39, Arg40, Tyr254, and Arg309 of NuoH, as well as Glu58 of NuoB (Fig. 4F). At the present resolution it is not possible to tell if the redox state of MQ9 differs in these two different positions. However, it is worth noting that all the residues that interact with MQ in the semi-inserted position are capable of hydrogen bonding, which could suggest the more polar semiquinone or fully reduced quinol forms of MQ binds in this position.

## Discussion

Structurally, mycobacterial Complex I is similar to Complex I from other prokaryotes. However, the presence of a two-component response regulator protein as a subunit of Complex I is unique to mycobacteria. Several mycobacterial pathogens possess orthologues of *MSMEG_2064*, such as *Rv3143, MAV_4032*, and *Mb3167, MUL_2453* in *M. tuberculosis, M. avium, M. bovis*, and *M. ulcerans*, respectively. Interestingly, an orthologue of *MSMEG_2064* is not present in *M. leprae*, which lacks the *nuo* operon and only retains a *nuoN* pseudogene. Other supernumerary subunits in Complex I from other species have been found in positions similar to MSMEG_2064 in mycobacterial Complex I. In *E. coli* Complex I the binding site for MSMEG_2064 is unoccupied (33) but Nqo16 is found in this position in the *T. thermophilus* complex (32). Nqo16 is a *Thermus*-specific protein of unknown function that is not required for stability of the complex, redox activity, or proton pumping (48). In Complex I from mitochondria, the supernumerary subunit NDUFA9 occupies a position equivalent to MSMEG_2064 (26, 28). NDUFA9 has been linked to Complex I assembly and stabilization of the connection between the peripheral arm and membrane region and has been proposed to have a redox sensing activity (83). However, unlike MSMEG_2064, neither Nqo16 in *T. thermophilus* or NDUFA9 in mitochondria have been linked to signalling. Therefore, if MSMEG_2064 has two-component response activity, its role appears to be mycobacteria-specific. Prediction of MSMEG_2064 interaction partners using STRING (84) with a high confidence cutoff (0.700) identified NuoE, NuoC, NuoG, of which the interaction between MSMEG_2064 and NuoC is consistent with the structure presented here. In addition, this analysis predicted interaction of MSMEG_2064 with MSMEG_2248 and MSMEG_0980, two predicted kinases of unknown function. Based on the structure, phosphorylation of MSMEG_2064 could modulate ETC activity by preventing its association with Complex I, which would in turn prevent Complex I assembly.

The GTP-like moiety bound in mycobacterial NuoG distinguishes it from homologues in other species. This additional cofactor may simply serve a structural role, as has been proposed for cluster N7 (36). However, it is possible the GTP moiety of NuoG serves as the precursor to a molybdopterin that catalyzes a yet to be discovered reaction in mycobacterial Complex I. Alternatively, the proximity of this density to an iron-sulfur cluster may suggest a redox sensing function, similar to the redox-inactive NADPH cofactor in the NDUFA9 subunit of eukaryotic Complex I (83). Interestingly, in the molybdopterin cytosine dinucleotide (Mo-CDP)-containing carbon monoxide dehydrogenase from the aerobic bacterium *Hydrogenophaga pseudoflava*, culture in the absence of molybdenum results in the replacement of Mo-CDP with a cytosine nucleotide and a functionally inactive enzyme (85). NuoG has been implicated in immune evasion by *M. tuberculosis* (86), and deletion of the *nuoG* gene in the tuberculosis vaccine candidate *M. bovis* BCG ΔureC::hly was found to increase its protection compared to the parental BCG ΔureC::hly candidate (87). The GTP-like moiety detected in Complex I was from mycobacteria grown in laboratory conditions, which only partially mimic the biological niche of mycobacteria. Therefore, the role of the purine triphosphate moiety, which appears unique to mycobacterial Complex I, requires further study in mycobacteria from more complex environments.

Plasticity in the NADH dehydrogenase system (24) is a notable attribute of the highly branched and flexible mycobacterial ETC (1, 9). The results presented here are consistent with this plasticity occurring, at least in part, due to expression of Complex I, which can range from high expression in carbon-limited aerobic conditions, to low but detectable in carbon-abundant conditions, to undetectable in hypoxic conditions. Downregulation of mycobacterial Complex I in hypoxic conditions has been proposed to decrease the contribution of the ETC to the PMF, with the electrogenic export of succinate used for maintaining the PMF in the absence of oxygen as a terminal acceptor (23, 88). We did not observe a growth defect in a *ΔnuoF* strain, but this lack of observed phenotype could be due to Ndh providing sufficient NADH dehydrogenase activity for growth in the culture conditions that were used. Complex I activity is a major contributor to cellular reactive oxygen species (ROS) (89) and Complex I expression is downregulated in oocytes to reduce ROS production and preserve cellular fitness (90). A similar downregulation of mycobacterial Complex I could support persistence in hypoxic conditions. The presence of the orphan two component response regulator protein MSMEG_2064 and a GTP-like moiety in the NuoG subunit hint at additional roles for Complex I in the sophisticated and flexible ETC of mycobacteria.

## Author contributions

YL prepared *M. smegmatis* 3×FLAG tagged Complex I strains and the *MSMEG_2064* deletion strain, cultured bacteria, purified protein, and performed electron microscopy and image analysis. AP performed mass spectrometry experiments. SAB prepared the *M. smegmatis* 3×FLAG tagged Ndh strain and the *ΔnuoF* deletion strain and characterized growth of the *ΔnuoF* strain. JL provided guidance on *M. smegmatis* molecular genetics. PB advised on interpretation of the structures. SV designed and supervised mass spectrometry experiments. JLR supervised the research and coordinated experiments. YL and JLR wrote the manuscript and prepared figures with input from the other authors.

## Acknowledgements

YL was supported by a Canadian Institutes of Health Research Canada Graduate Scholarship and a SickKids Research Institute Restracomp Scholarship. AP was supported by an Ontario Graduate Scholarship. This research was funded by Canadian Institutes of Health Research grants PJT162186 (JLR), PJT451412 (SV), PJT173353 (JL). JLR was supported by the Canada Research Chairs program. CryoEM data were collected at the Toronto High-Resolution High-Throughput Cryo-EM facility, supported by the Canada Foundation for Innovation and Ontario Research Fund.

## Data availability

Electron microscopy maps are available from the electron microscopy databank with accession codes EMD-27963, EMD-27964, and EMD-27965 and atomic models are available from the protein databank with accession codes 8E9G, 8E9H, and 8E9I. Mass spectrometry data are available from the massive database as entry MSV000090235.

## Materials and Methods

### Generation of strains

*M. smegmatis* strains were generated with the ORBIT method (91). ORBIT requires transformation of the starting bacterial strain (*M. smegmatis* mc^2^155) with pKM444 (Addgene #108319), a plasmid that encodes a Che9c phage RecT annealase and a Bxb1 integrase. To generate strains encoding a 3×FLAG sequence 3’ of the reading frame of the gene of interest, the pKM444 transformed strain was subsequently transformed with an oligonucleotide that guides integration of the payload plasmid pSAB41, a modification of pKM491 (Addgene #109282) described previously (92). Incorporation of pSAB41 into the *M. smegmatis* genome results in fusion of a 3×FLAG tag at the C terminus of the protein of interest. Transformants were selected with 50 μg/mL Hygromycin B and proper insertion was confirmed by colony PCR.

To generate strains containing gene deletions, pKM444-transformed *M. smegmatis* mc^2^155 was subsequently transformed with the targeting oligonucleotide and the payload plasmid pKM496 (Addgene #109301), which replaces the gene of interest with a Zeocin (phleomycin D1) resistance gene. The *nuoF* knockout strain was prepared from a pKM444 background strain. Knockout of the *MSMEG_2064* gene was prepared from the *M. smegmatis* 3×FLAG NuoH strain. Transformants were selected with 25 μg/mL Zeocin and proper insertion was confirmed by colony PCR. The oligonucleotide sequences used for the generation of all strains is provided in the supplementary information (Table S5).

### Bacterial culture and membrane harvest

*M. smegmatis* was cultured in batches of 1 L in Fernbach flasks (total flask volume 2.8 L) containing Middlebrook 7H9 broth supplemented with 0.05% (v/v) tween-80 and one of: albumin (0.5% w/v), dextrose (11 mM or 5.5 mM), glycerol (11 mM), succinate (11 mM), or ADS (0.5% [w/v] albumin, 11 mM dextrose, and 14 mM NaCl). A culture containing only 7H9 supplemented with 0.05% (v/v) tween-80 was also prepared. Cultures were grown at 30 °C in a shaking incubator at 180 rpm for 72 h before harvesting membranes. To compare Complex I expression under different oxygenation conditions, 1.9 L cultures of 7H9 supplemented with 11 mM dextrose, 5.5 mM dextrose, or 11 mM succinate with a headspace ratio of 0.5 following Wayne’s model of hypoxia (43) were prepared. Flasks were sealed with plastic food wrap. Hypoxic cultures were incubated at 30 °C with 120 rpm shaking for 72 h before membrane harvest. Large-scale aerobic cultures for protein purification were grown in 7H9 supplemented with 11 mM succinate and 0.05% (v/v) tween-80 at 30 °C and 180 rpm.

Cells were collected by centrifugation at 6500 *g*. To harvest membranes for western blotting, cells from each culture condition (1 L or 1.9 L) were resuspended in 20 to 25 mL PBS (80 mM Na_2_HPO_4_, 20 mM NaH_2_PO_4_, 100 mM NaCl, pH 7.2 at 25 °C). Cells were lysed by sonication on ice (10 cycles of 20 s sonication with 10 s pause, at 25% amplitude) with a Q500 probe sonicator (QSonica). Cell debris was removed by centrifugation at 33,745 *g* for 30 min and membranes were collected by centrifugation at 200,000 *g* for 60 min. To harvest membranes for protein purification, resuspended cells were lysed with four passages at >20 kpsi with an EmulsiFlex-C3 High Pressure Homogenizer (Avestin). Cells from 1 L of culture were resuspended in 20 mL of lysis buffer (50 mM MES pH 6.0 at 4 °C, 100 mM NaCl, 5 mM benzamidine HCl, 5 mM aminocaproic acid, and 1 mM PMSF). Unbroken cells and debris were removed by centrifugation at 39,000 *g* for 30 min. Membranes were collected from the supernatant by centrifugation at 200,000 *g* for 60 min. Membranes were resuspended in buffer (20 mM MES pH 6.0 at 4 °C, 15% [v/v] glycerol, 5 mM CaCl_2_, 100 mM NaCl, and 1 mM PMSF) before storage at −80 °C.

### Western blotting

Membranes were solubilized in 500 to 1000 μL PBS with 2% (w/v) sodium dodecyl sulfate (SDS) for 2 h at 50 °C with occasional mixing. Total protein content in the solubilized membranes was quantified by bicinchoninic acid (BCA) assay (ThermoFisher Scientific). Protein samples (4 μg) was loaded on a 15% SDS gel in duplicate. Following gel electrophoresis, proteins were either transferred onto 0.45 μm nitrocellulose membrane (Bio-Rad) for western blotting or stained with Coomassie Brilliant Blue (Bio-Rad) to ensure consistent loading. M2 anti-FLAG primary antibody (Sigma) at a 1:5000 dilution was used for detection of the 3×FLAG tag. Horseradish peroxidase-conjugated anti-mouse IgG (Bio-Rad) at a 1:3000 dilution was used as the secondary antibody. All antibody dilutions were made in TBS-T (50 mM Tris pH 7.6 at 25 °C, 150 mM NaCl, 0.1% [v/v] Tween-20, 5% [w/v] bovine serum albumin). Bands were visualized using SuperSignal West Pico Chemiluminescent Substrate (ThermoFisher Scientific) with a ChemiDoc XRS+ imaging system (Bio-Rad).

### Protein purification and activity assay

All purification steps were conducted at 4 °C. Frozen membranes were thawed and solubilized by gentle stirring with 1% (w/v) DDM. Insoluble material was removed by centrifugation at 200,000 *g* for 50 min followed by filtration with a 0.45 μm syringe filter (Millipore). Solubilized protein was loaded onto 1.5 mL of M2 affinity matrix (Sigma) equilibrated with wash buffer (20 mM MES pH 6.0, 15% [v/v] glycerol, 5 mM CaCl_2_, 100 mM NaCl, and 0.03% [w/v] DDM). Complex I was eluted with 4.5 mL of 150 μg/mL 3×FLAG peptide in wash buffer. The sample was concentrated and loaded onto a Superose 6 Increase 10/300 GL size exclusion column (Cytiva) equilibrated in wash buffer. Eluted Complex I fractions were pooled and concentrated and protein concentration was measured by BCA assay. Assay of NADH:quinone oxidoreductase activity was conducted at 37 °C with a BioTek Synergy Neo2 plate reader (Agilent) in buffer composed of 10 mM MES pH 6.0, 25 mM NaCl, 5 mM CaCl_2_, 0.05% 3-[(3-cholamidopropyl)dimethylammonio]-1-propanesulfonate (CHAPS), and 0.05% asolectin. The concentration of NADH in the assay mixture was followed spectrophotometrically by measuring absorbance at 340 nm. To calibrate absorbance at 340 nm with NADH concentration, an NADH standard curve was prepared in the same buffer. Complex I (1 μg in 200 μL reaction volume) was incubated with 200 μM of either menadione or decylubiquinone for 10 min before starting the reaction with the addition of 200 μM NADH. To examine inhibition of Complex I, either 50 μM or 150 μM of rotenone was added prior to the 10 min incubation.

### Culture in presence of molybdenum and formate dehydrogenase activity assay

*M. smegmatis* was cultured in 7H9-succinate (11 mM) media supplemented with 10 μM ammonium molybdate. Complex I (3×FLAG NuoH) was purified and quantified as described above. Assay of formate:NAD^+^ oxidoreductase activity was conducted as described in the previous section at 37 °C with a BioTek Synergy Neo2 plate reader (Agilent) in buffer composed of 10 mM MES pH 6.0, 25 mM NaCl, 5 mM CaCl_2_, 0.05% (w/v) 3-[(3-cholamidopropyl)dimethylammonio]-1-propanesulfonate (CHAPS), and 0.05% (w/v) asolectin. Complex I (1 μg) or *Candida boidinii* formate dehydrogenase (0.015 U, Megazyme) was incubated with 30 mM of sodium formate for 5 min before starting the reaction with the addition of 200 μM of NAD^+^ in a total reaction volume of 200 μL. NADH formation was monitored spectrophotometrically by measuring absorbance at 340 nm.

### Trypsin digestion for mass spectrometry

Mass spectrometry analysis was conducted on purified Complex I samples following trypsin digestion both in solution and following separation of proteins by gel electrophoresis. For in solution trypsin digestion *M. smegmatis* Complex I (8 μg) was digested with MS-grade trypsin/Lys-C mix (Promega) at a 25:1 (w/w Complex I:protease ratio in 25 mM Tris-HCl, pH 8.0) at 37 °C for 16-18 hours. The reaction was quenched with the addition of trifluoroacetic acid (TFA) to a final concentration of 0.5% (v/v). Digested samples were flash frozen in liquid nitrogen and stored at −80 °C until LC-MS analysis. For in gel trypsin digestion, bands were excised from a 15% SDS polyacrylamide gel following Coomassie blue staining. Excised bands were destained with 25 mM NH_4_HCO_3_ (pH 7.0) and 50% (v/v) acetonitrile and dehydrated in a refrigerated vacuum concentrator (Speedvac - Thermo Fisher). MS-grade trypsin/Lys-C mix (200 ng, Promega) in 25 mM Tris-HCl (pH 8.0) was added to the dehydrated gel fragments and the sample was digested overnight at 37 °C following addition of 50 mM NH_4_HCO_3_ (pH 7.0). Digested peptides were extracted through sonication in 5% (v/v) formic acid and 50% (v/v) acetonitrile, dried by refrigerated vacuum concentrator, and stored at −20 °C. Prior to mass spectrometry analysis, samples were resuspended in 0.1% formic acid.

### LC-MS/MS

Reverse phase separation of the digests was conducted at 40.0 °C with a bridged ethylene hybrid C18 (1.7 μm particle size; 1 mm ×100 mm) ACQUITY column (Waters). Peptides were separated on a 40 min H_2_O:acetonitrile gradient at flow rate of 0.150 mL/min, by linearly ramping the acetonitrile content from 2% to 50%. Both mobile phases were acidified with 0.1% (v/v) formic acid. In-gel digested samples (5 μL) and in-solution digested samples (0.2-2.0 μg) were injected. Eluents were directed to a Synapt G2Si quadrupole time-of-flight mass spectrometer (Waters) equipped with a standard electrospray ionization source operated in positive ion mode at +3 kV. The desolvation temperature was 600 °C and the cone and source offset voltages were 40 and 50 V, respectively. Peptides and proteins present in the sample were identified with both data-dependent acquisition (DDA) (93) and data-independent acquisition (DIA) (94) experiments. For both DIA and DDA survey scans, mass spectra were recorded within the m/z range of 50-2000 Th with a scan time of 0.4 seconds. For both the DIA and the survey scans of DDA experiments, the quadrupole was operated in RF-only mode with a manual profile to excludes the transmission of ions with m/z values lower than 240 Th (95). The raw DIA and DDA data were searched against the Uniprot *M. smegmatis* (strain ATCC 700084/ mc^2^155) database containing 12,794 proteins (downloaded on 2021-Oct-18), with ProteinLynx Global Server v. 3.5.3 (Waters). Table S4 includes all the identified proteins and peptides. All raw data are available for download from the MassIVE databank (entry #MSV000090235).

### CryoEM sample preparation and data collection

Holey gold grids with 2 μm holes arranged in a square lattice were nanofabricated (96). Purified Complex I was concentrated to ~2 mg/mL following size exclusion chromatography and glycerol was removed with a Zeba Spin desalting column (ThermoFisher Scientific). Grids were glow-discharged for 2 min in air immediately prior to application of 2 μL of sample to each grid. Grids were then blotted in an EM GP2 Plunge Freezer (Leica Microsystems) for 0.5 s before freezing in liquid ethane.

For sample optimization, specimens were screened with a FEI Tecnai F20 electron microscope operating at 200 kV and equipped with a Gatan K2 Summit direct electron detector. Images were collected at 25,000× magnification in counting mode, corresponding to a pixel size of 1.45 Å at an exposure rate of ~5 electrons/pixel/s. High-resolution data was collected with a Titan Krios G3 electron microscope (ThermoFisher Scientific) operating at 300 kV and equipped with a Falcon 4i direct electron detector camera (ThermoFisher Scientific). Data collection was automated with the EPU software (ThermoFisher Scientific). A total of 4,002 movies, each consisting of 30 exposure fractions, were collected at a nominal magnification of 75,000× corresponding to a calibrated pixel size of 1.03 Å with an exposure rate of ~6 e^−^/pixel/s and a total exposure of ~47 e^−^/Å^2^.

### CryoEM image analysis

Assessment of data quality during automated collection was conducted with cryoSPARC Live (97). Unless otherwise stated, image analysis was conducted with cryoSPARC v3.3.1 (98). Exposure fractions were aligned with MotionCor2 (99) using a 7×7 grid. Contrast transfer function (CTF) parameters were estimated in patches. Templates for particle picking were generated from projections of a lower resolution 3D map. Selected particle images were extracted with a 440×440 pixel box yielding 357,064 particle images. The dataset was cleaned with several rounds of ab-initio reconstruction and heterogeneous refinement to yield 63,684 particle images. Image parameters were then converted to RELION (100) .star file format with pyem (https://doi.org/10.5281/zenodo.3576630) and individual particle motion-correction was performed with Bayesian Polishing (101), with the pixel size binned to 1.18 Å. Next, particle images were reimported to cryoSPARC, per-particle CTF parameters re-estimated, and the dataset was cleaned again with several rounds of ab-initio reconstruction and heterogeneous refinement to yield 60,177 particle images. 3D variability analysis with a mask around the quinone binding channel allowed for separation of the dataset into 14,835 and 45,342 particle images corresponding to the semi-inserted and fully-inserted quinone binding positions respectively. Final 3D maps were calculated with non-uniform refinement (102).

### Model building and refinement

Starting models of NuoA–NuoN were generated by one-to-one threading with the Phyre2 server (103) using the *M. smegmatis* amino acid sequences and predicted models of *M. tuberculosis* subunits from the AlphaFold Protein Structure Database (104). The starting model for MSMEG_2064 was generated with AlphaFold-Colab (105). Models were rigid-body fitted into the consensus cryoEM map in Chimera with the *T. thermophilus* Complex I structure (PDB: 4HEA) as a guide. Optimization of model-to-map fit and atomic model dihedral angles was done with Coot v0.9.6 (106), followed by several rounds of refinement with ISOLDE v1.3 (107) and PHENIX v1.19.2 (108). The consensus model was then rigid-body-fitted into the maps showing semi-inserted and fully-inserted quinone and optimization and refinement were conducted as in the consensus model. For AcPIM_2_, an atomic model was created in ChemDraw v20.1 (PerkinElmer) and the 3D model was generated with *phenix*.*elbow* from the SMILES string of the atomic model. Fitting of AcPIM_2_ in the cryoEM density was done in Coot v0.9.6 as with fitting of other ligands. Figures were rendered with UCSF Chimera (109) and UCSF ChimeraX (110).

## Figures

**Supplementary Figure 1:**
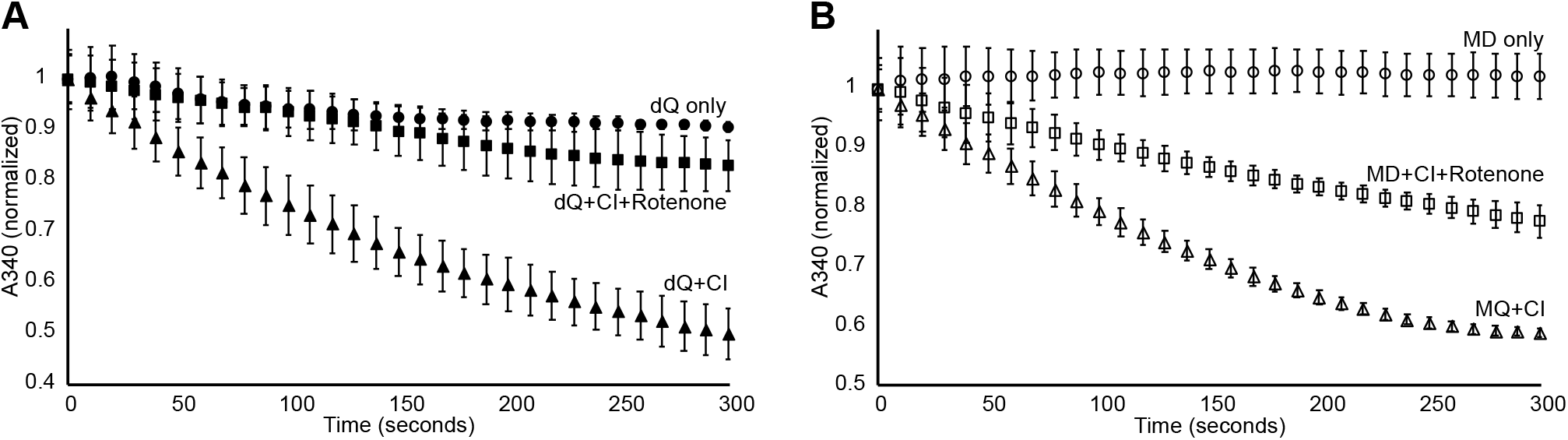
Replicate of NADH:quinone oxidoreductase assay from independent protein purification. **A**, Complex I NADH:decylubiquinone (dQ) oxidoreductase activity with and without 150 mM rotenone. **B**, Complex I NADH:menadione (MD) oxidoreductase activity with and without 150 mM rotenone.

**Supplementary Figure 2:**
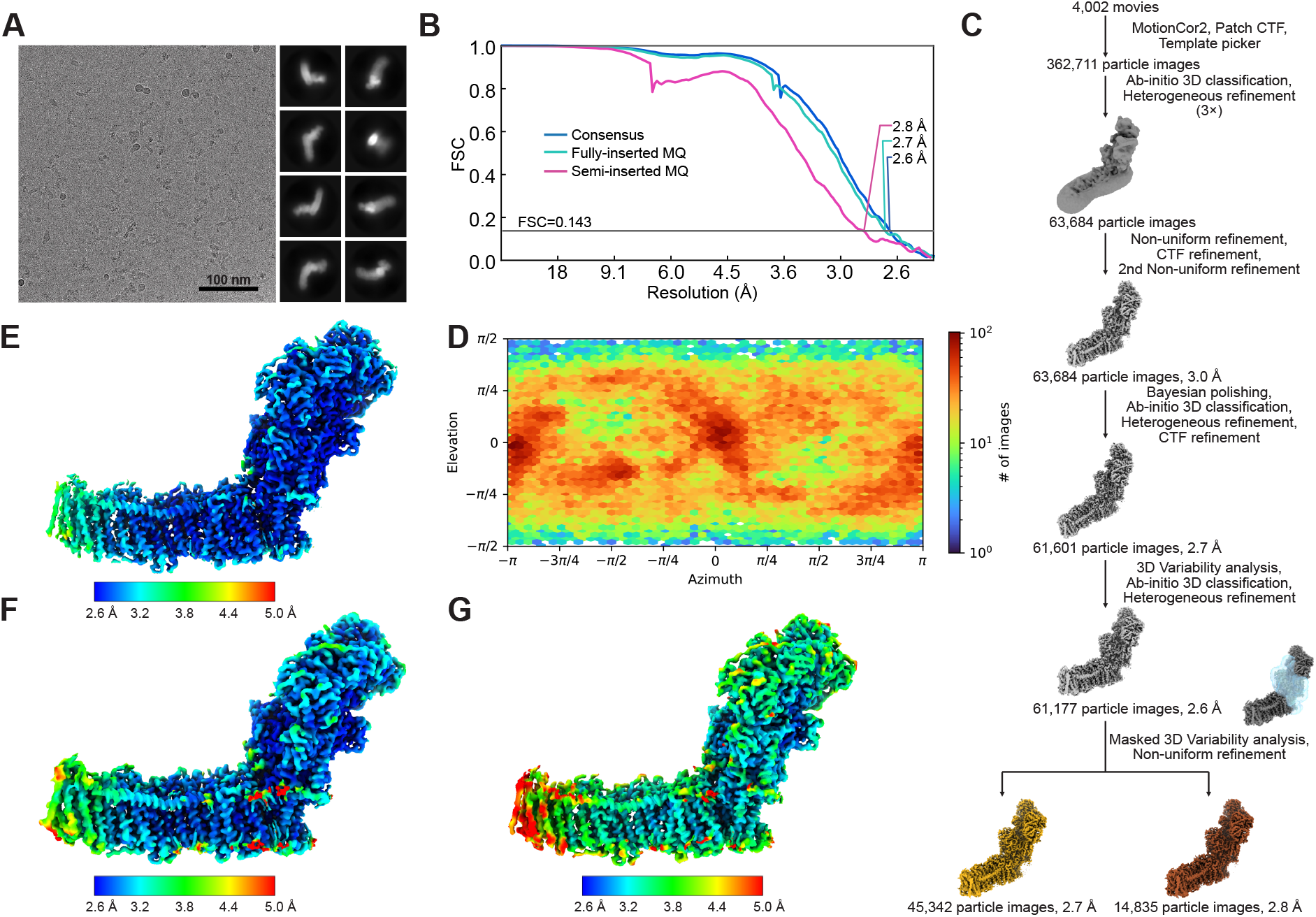
CryoEM of mycobacterial Complex I. **A**, Representative cryoEM micrograph (*left*) and representative 2D class averages (*right*). **B**, Fourier Shell Correlation (FSC) curve of the consensus map (*blue*), the fully-inserted quinone map (*cyan*), and the semi-inserted quinone map (*magenta*) following gold standard refinement and correction for masking. **C**, CryoEM data processing workflow. **D**, Viewing direction distribution plot of particle images. **E**, Local resolution in the consensus map. **F**, Local resolution of the fully-inserted quinone map. **G**, Local resolution in the semi-inserted quinone map.

**Supplementary Figure 3:**
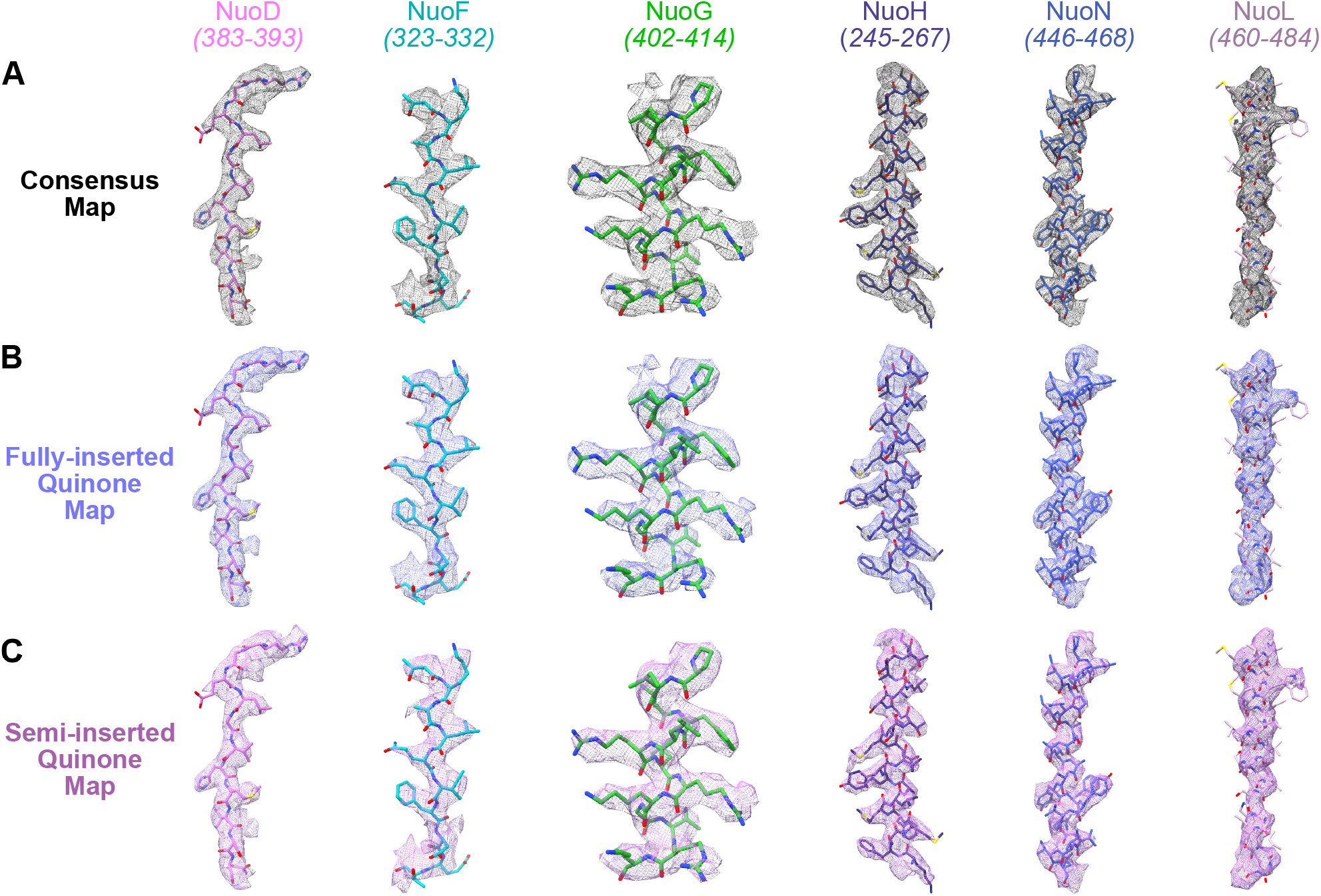
Examples of model-in-map fit. Examples are shown to illustrate the variable resolution of the cryoEM map from the consensus map (**A**), fully-inserted quinone map (**B**), and semi-inserted quinone map (**C**).

**Supplementary Figure 4:**
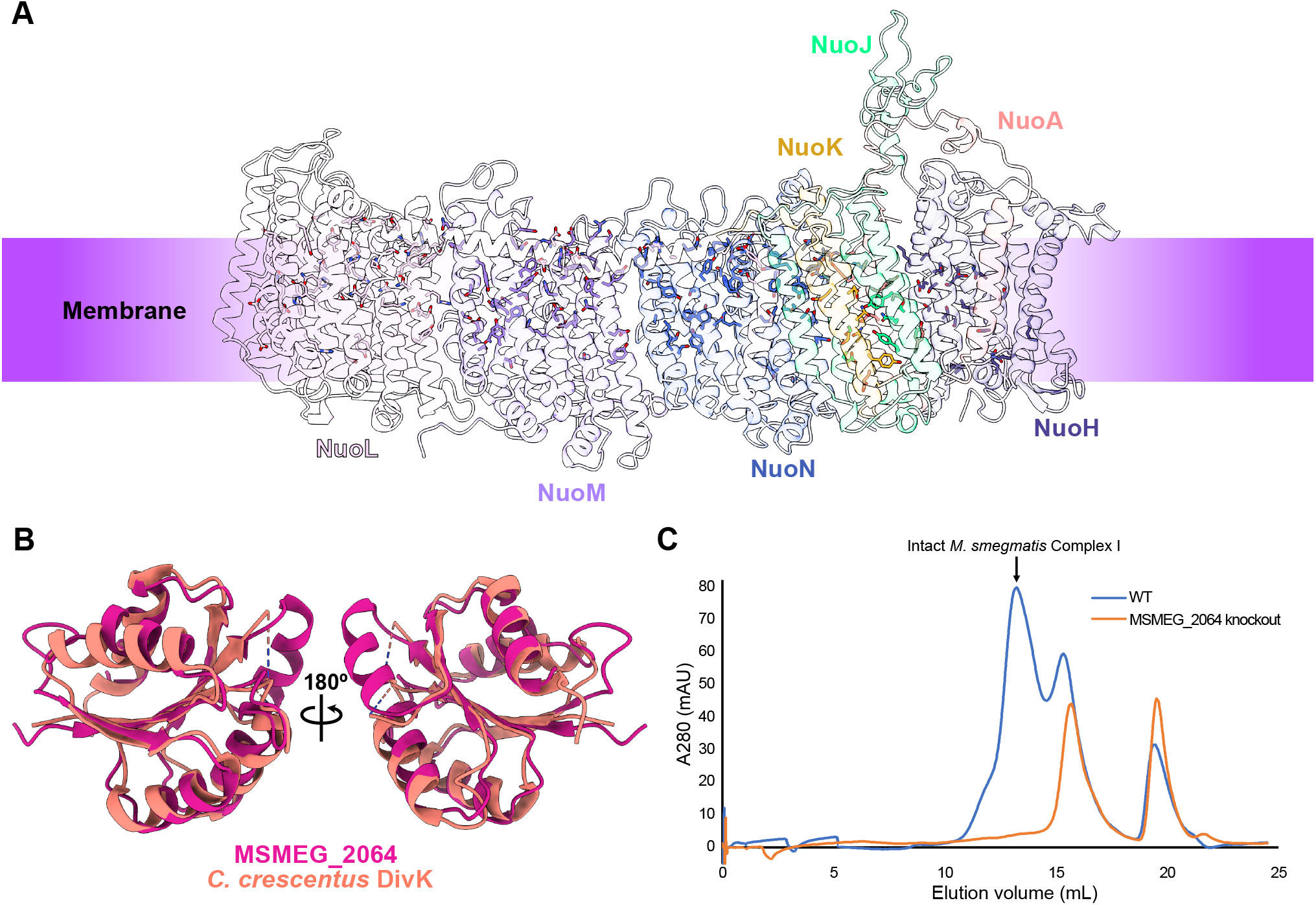
Membrane subunits and MSMEG_2064. **A**, Distribution of polar (Tyr, Ser, Gln, Asn) and charged (Asp, Glu, Lys, Arg, His) residues in the membrane region. **B**, Alignment of the atomic model of MSMEG_2064 with *C. crescentus* DivK (PDB: 1M5U). **C**, Gel filtration chromatogram overlay of 3×FLAG-tagged Complex I purified from a wild-type background (*blue curve*) and 3×FLAG-tagged Complex I purified from a MSMEG_2064 knockout background (*orange curve*). The peak corresponding to intact Complex I is indicated with an arrow.

**Supplementary Figure 5:**
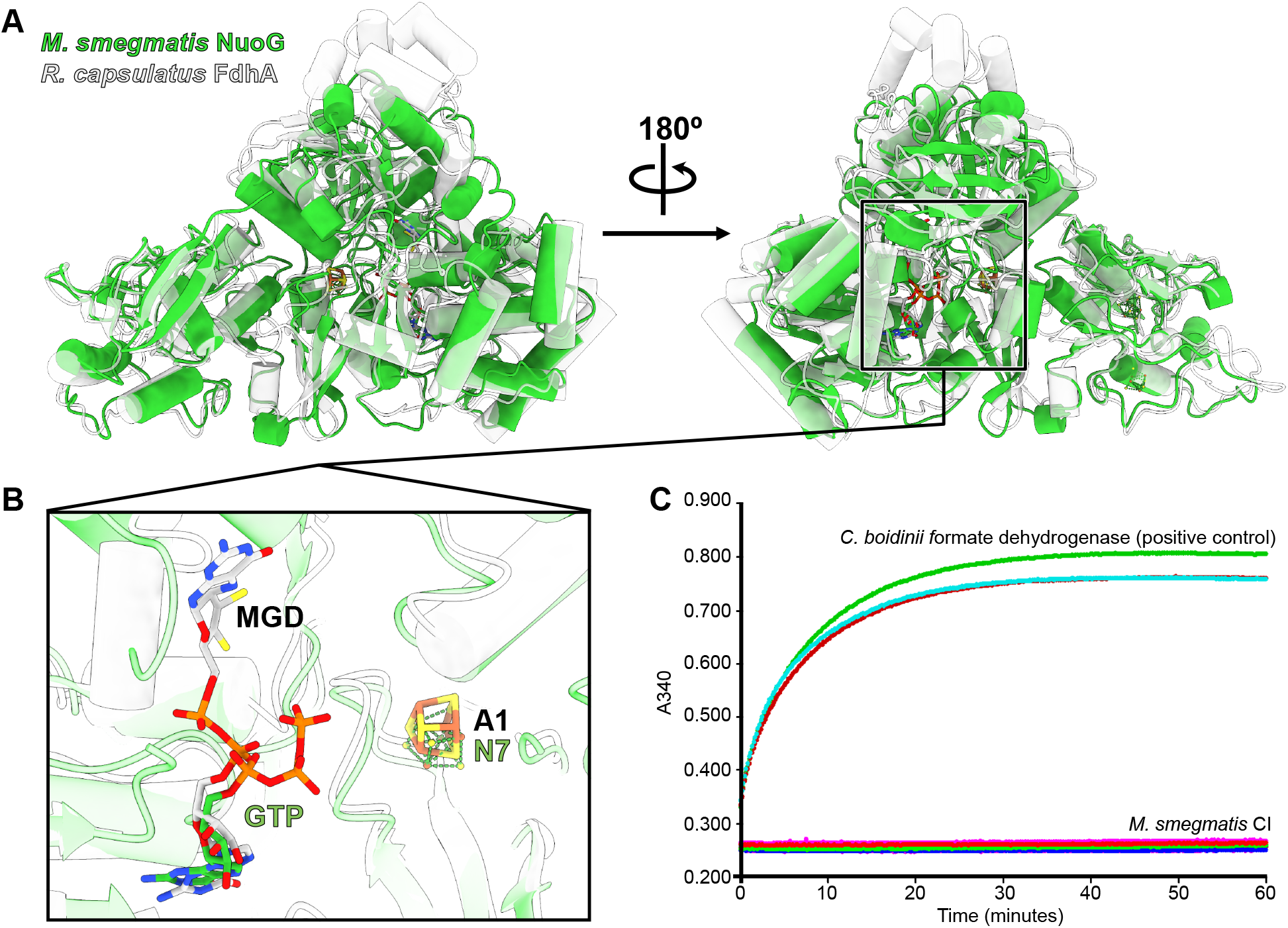
Comparison of *M. smegmatis* NuoG with formate dehydrogenases. **A**, Alignment of the atomic model of *M. smegmatis* NuoG (*lime green*) with *R. capsulatus* FdhA (PDB: 6TG9, *light gray*). Location of the GTP moiety/Mo-bisMGD and the proximal 4Fe–4S cluster are indicated by the square. **B**, Closeup view of the alignment of cofactors between *M. smegmatis* NuoG (*lime green*) and *R. capsulatus* FdhA (*light gray*). **C**, Assay of formate:NAD^+^ oxidoreductase activity of molybdenum-supplemented *M. smegmatis* Complex I compared to formate dehydrogenase from *C. boidinii*.

**Supplementary Figure 6:**
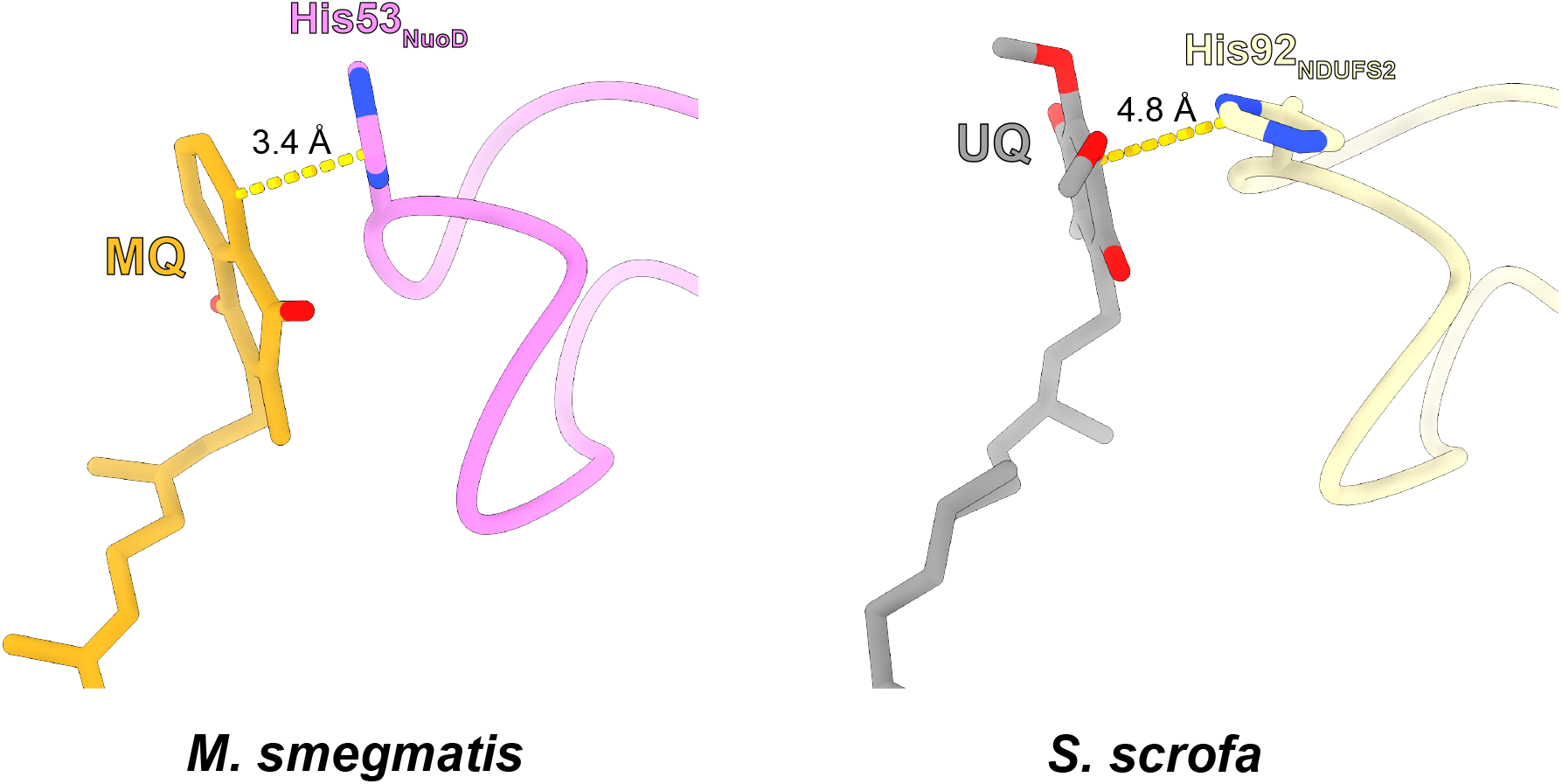
MQ-His and UQ-His interaction distances. Comparison of the distance between MQ and His53 of NuoB from *M. smegmatis* and the position between UQ and His92 of NDUFS2 from *S. scrofa* (PDB: 7V2C). **Video 1: 3DVA component 1** **Video 2: 3DVA component 2**

**Table S1:**
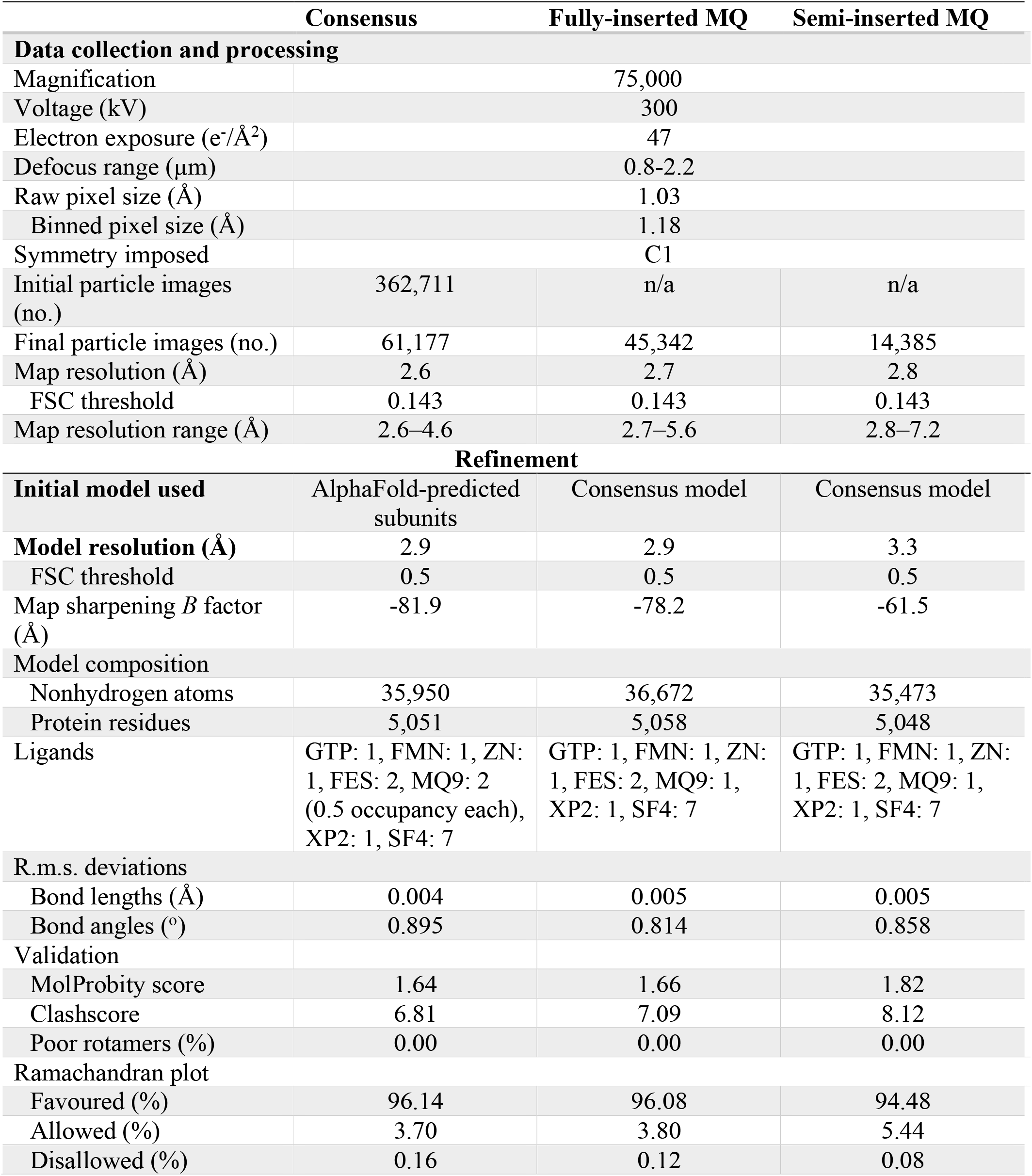
CryoEM and atomic model building statistics

**Table S2:** Chain IDs and number of residues built

| Subunit name | Chain | Total number of residues | Residues built |  |  |
| --- | --- | --- | --- | --- | --- |
|  |  |  | Consensus | Fully-inserted MQ | Semi-inserted MQ |
| **MSMEG_2064** | O | 132 | 4–128 | 4–128 | 5–128 |
| **NuoA** | A | 122 | 1–122 | 1–122 | 2–122 |
| **NuoB** | B | 184 | 2–184 | 2–184 | 2–184 |
| **NuoC** | C | 238 | 19–238 | 19–238 | 19–237 |
| **NuoD** | D | 442 | 37–442 | 36–442 | 36–442 |
| **NuoE** | E | 245 | 6–238 | 6–237 | 6–237 |
| **NuoF** | F | 443 | 2–436 | 2–437 | 2–436 |
| **NuoG** | G | 794 | 13–792 | 12–793 | 13–792 |
| **NuoH** | H | 408 | 1–395 (375–395 register unconfirmed) | 1–395 (375–395 register unconfirmed) | 1–395 (375–395 register unconfirmed) |
| **NuoI** | I | 180 | 4–168 | 4–168 | 3–168 |
| **NuoJ** | J | 252 | 13–240 | 13–243 | 13–242 |
| **NuoK** | K | 99 | 1–99 | 1–99 | 1–99 |
| **NuoL** | L | 629 | 5–629 | 5–629 | 7–629 |
| **NuoM** | M | 529 | 4–519 | 4–520 | 4–521 |
| **NuoN** | N | 521 | 3–521 | 3–521 | 3–520 |

**Table S3:**
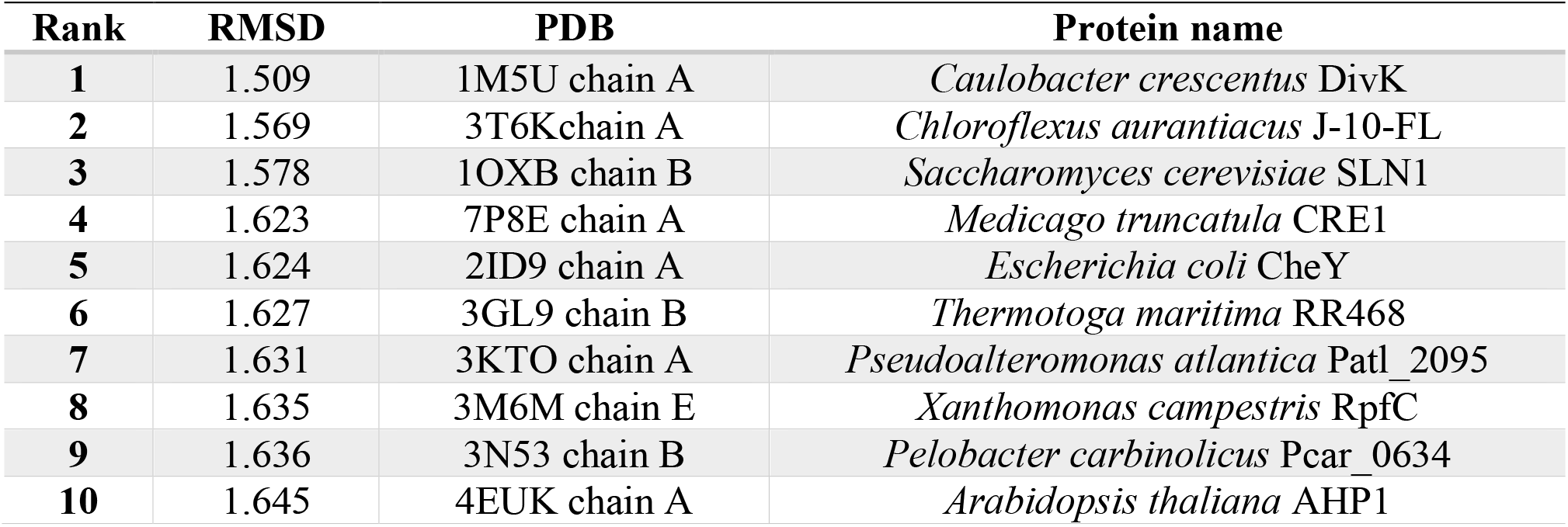
Top ten PDB entries with lowest Cα RMSD compared to MSMEG_2064 (consensus model chain A) by PDBeFold as of July 2022

**Table S4: Mass spectra of purified *M. smegmatis* Complex I (.xlsx format)**

**Table S5:**
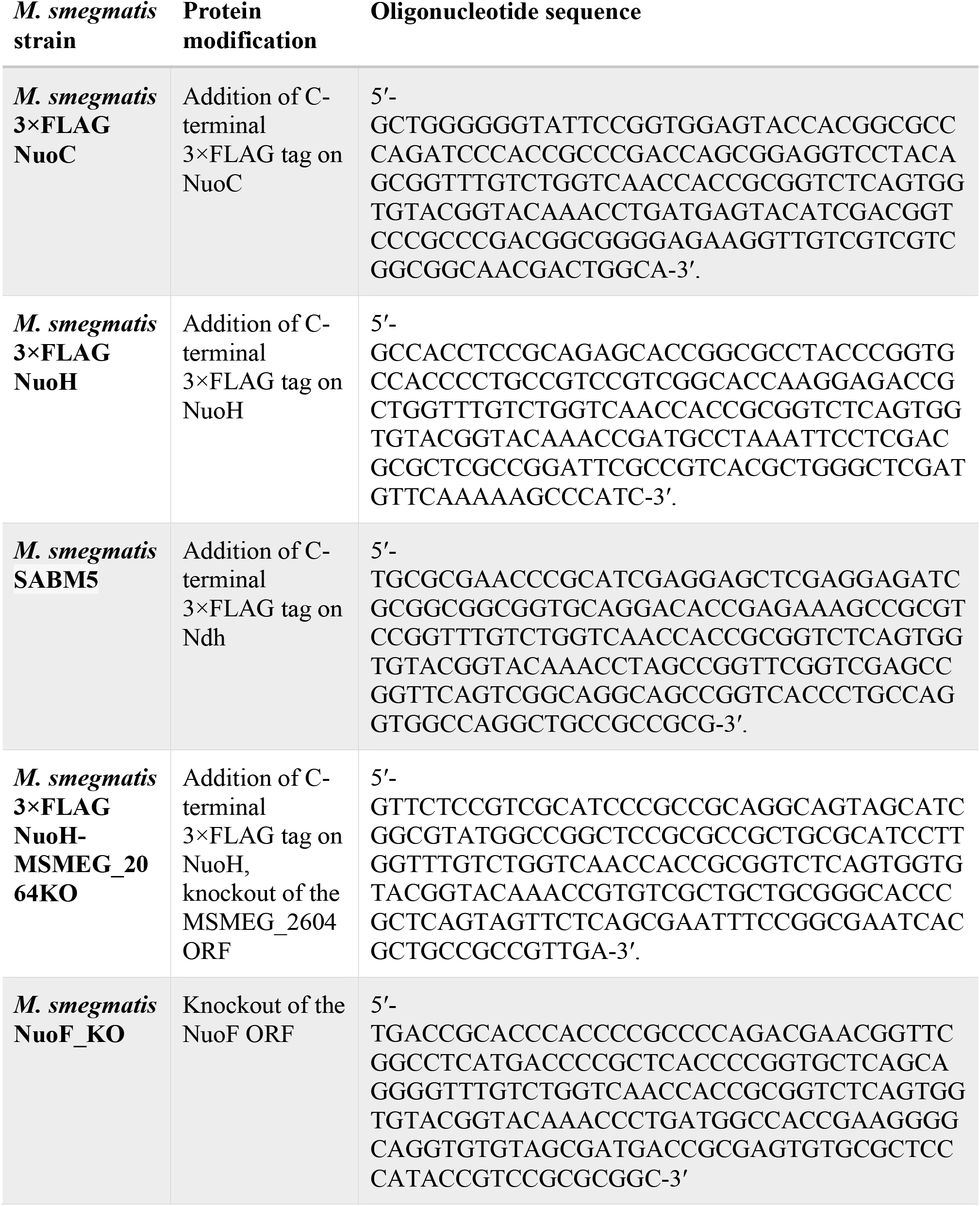
ORBIT oligonucleotide sequences

